# Amplification of genomic regions harbouring genes with dose-limiting effects provides selection pressure for acquiring aberrant epigenetic silencing in ovarian cancer

**DOI:** 10.1101/630228

**Authors:** Nair Bonito, Ieva Eringyte, Nahal Masrour, Charlotte Wilhelm-Benartzi, Robert Brown, Edward Curry

## Abstract

High-grade serous ovarian cancer (HGSOC) feature widespread genomic rearrangements that alter the copy number of genes. Genes for which elevated expression is detrimental to growth may be passengers in rearrangements that increase the copy number of strongly selective driver genes. There would then be selection pressure for compensatory epigenetic silencing of such passenger genes, such as through promoter DNA methylation.

We have used two independent cohorts of primary HGSOC tumour samples to provide evidence of consistent dosage-compensating promoter methylation (DCPM) genome-wide. Mapping CpG methylation to genomic copy number, we show that bias due to variable tumour cellularity of tissue samples results in false positive associations between methylation and copy number. Adjusting for this bias, we found that approximately 5-10% of all measured promoter CpGs with copy number gain showed a statistically significant increase in methylation. DCPM nullifies the association between increased copy number and increased gene expression. We confirmed a functional basis for selective pressure against over-expressing two candidate passenger genes in the 3q12 locus, as overexpressing either gene reduced spheroid formation efficiency in two HGSOC cell lines. DCPM represents a novel class of specific functional interactions between genetic and epigenetic landscapes of cancers.

## INTRODUCTION

Epithelial ovarian cancer is the most lethal gynaecological cancer, with only 46% of patients estimated to survive beyond the 5-year mark. This can be attributed to late diagnosis in most cases, at which stage the cancer has spread, metastasising to the peritoneal cavity (Lengyel 2010). Although ovarian cancer is commonly grouped as a single disease, subtypes exhibit significant phenotypic and molecular differences (Vaughan et al. 2011).

High-grade serous ovarian cancer (HGSOC) is characterised by near-ubiquitous mutation of TP53 resulting in a loss of genomic integrity (Consortium 2011a). This genetic instability leads to widespread changes in genomic copy number, as a consequence of structural rearrangements. Such copy-number variations (CNVs) have been widely studied in cancers (Bergamaschi et al. 2006; Beroukhim et al. 2010). This has led to the identification of a number of cancer driver and tumour-suppressor genes, characterized by recurrent amplification or deletion. Changes in gene dosage as a consequence of genomic CNVs typically result in corresponding changes in gene expression (McCarroll et al. 2006). In fact, one popular approach to distinguish oncogenic drivers from ‘passengers’ within a genomic region of CNV is by looking for genes with the greatest and most consistent change in gene expression in accordance with the CNV status (e.g. (Akavia et al. 2010; Beroukhim et al. 2010)). In a landscape study profiling genomic changes in HGSOC, 113 recurrent CNV events were identified (Consortium 2011a). This included 30 aberrations affecting extended genomic regions: 5 gains and 18 losses each occurred in more than half of the 489 tumours profiled. 113 recurrent focal CNVs were also identified, including amplifications of CCNE1, MYC and MECOM, and deletions of PTEN, RB1 and NF1. In contrast, the same study only identified 9 genes affected by recurrent point mutations (TP53, BRCA1, BRCA2, RB1, NF1, FAT3, CSMD3, GABRA6 and CDK12), which reinforces the view that the molecular pathology of HGSOC is predominantly driven by CNVs and not point mutations (Ciriello et al. 2013). Additionally, many cancers harbour aberrant epigenetic features: such as reversible chemical modifications on DNA or histone tails that can alter gene expression (Baylin and Jones 2011). Promoter DNA methylation in HGSOC has been found to silence expression of tumour suppressor genes (Sellar et al. 2003; Consortium 2011a) and mediators of sensitivity to chemotherapy (Zeller et al. 2012; Bonito et al. 2016).

Epigenetic regulation of the potential impacts of gene dosage are familiar in terms of X-inactivation, where female mammalian cells feature widespread DNA methylation on one of their two copies of the X chromosome(Mohandas et al. 1981). Additionally, gene duplication events during evolution have been observed to be accompanied by DNA methylation(Chang and Liao 2011). Epigenetic dose compensation in cancer has been suggested in a number of recent studies, mostly in breast cancer cohorts. One study(Murria et al. 2015) found that tumours exhibiting higher promoter methylation across a panel of genes were more likely (relative to tumours with lower promoter methylation) to harbour copy-number aberrations across the same panel of genes. In another study featuring genome-wide analysis of 33 breast tumours, a significant increase in the average promoter methylation was found at genes affected by copy gain, which was hypothesised to function as a dosage compensating mechanism to reduce the expression level of amplified genes(Flanagan et al. 2010). Silencing of these genes was correlated with dedifferentiation and poor survival in breast cancer as well as involvement in signalling pathways commonly deregulated in cancer. An analysis of tumour samples from TCGA described ‘uncoupling’ of gene expression from copy-number in breast cancer based on the observation that the expression of some genes is not related to their copy number (Mohanty et al. 2017). This study reports a trend for genes with higher correlation between copy number and DNA methylation to have lower correlation between copy number and expression. However, there remains limited evidence in the literature of genome-wide analysis of the relationship between copy-number variation and aberrant DNA methylation in cancer, nor of attempts to explain the functional consequences of dosage-compensating epigenetic silencing.

In this study we sought to determine whether regions of CNV are universally associated with changes in DNA methylation, or if select loci are reliably associated with increased promoter methylation on CN gain in HGSOC. Such loci with clear dosage-compensating promoter methylation (DCPM) may define a novel class of dosage-dependent tumour-suppressors. A functional impact of dosage dependent gene expression of DCPM-affected passenger genes identified from primary tumour cohorts, on tumour formation and growth was characterized experimentally in 2 ovarian cancer cell lines.

## RESULTS

### Frequent DNA hypermethylation is observed at promoters within regions of genomic copy gain in high-grade serous ovarian cancer

Genome-wide profiles of copy-number, gene expression and DNA methylation were obtained for a cohort of 336 Primary HGSOC tissue samples from The Cancer Genome Atlas (TCGA) (Consortium 2011a), and for 78 Primary HGSOC tissue samples from the Australian Ovarian Cancer Study, available through the International Cancer Genome Consortium (ICGC) (Patch et al. 2015b). Summary statistics characterizing the CNV landscape in these cohorts are provided in Table 1. Patient characteristics are available in the original publications (Consortium 2011b; Patch et al. 2015b).

**Table 1.** Summary of CNV landscape in each cohort

| Cohort | Avg Amplicons Per Tumour | Avg #CpGs Amplified | Avg #Promoter CpGs Amplified | Avg #Genes Amplified |
| --- | --- | --- | --- | --- |
| ICGC | 44.81 | 22311.31 | 2215.19 | 884.91 |
| TCGA | 65.61 | 3155.62 | 1223.32 | 3231.21 |

We hypothesise that loci with strong positive correlations between copy number and methylation could indicate that a gene associated to such loci is one for which overexpression confers a disadvantage to growth of the tumour. To understand the general pattern of relationships between genomic copy number (gCN) and DNA methylation, we first sought to quantify the locus-by-locus associations between measurements of gCN and CpG methylation across the tumour samples in each cohort. As CpG methylation in promoter CpG islands (CGI) has been demonstrated as frequently leading to silencing of gene expression (Bird and Wolffe 1999), we have focused on CpG sites in promoter regions. Average methylation levels for the vast majority of these loci were confirmed to be low across both cohorts (Supplementary Figure S1). For both cohorts, we mapped DNA methylation level at each measured CpG site to its absolute genomic copy number estimate in each sample, then calculated the Spearman correlation coefficient relating the methylation β-values to copy number at the corresponding locus. It was clear that the observed statistical significance of positive Spearman correlations were dramatically greater (i.e. more correlated) than those expected if copy-number and DNA methylation were independent for all loci (Figure 1A-B). This was also true for loci with negative correlations between copy-number and DNA methylation (Supplementary Figure S2A-B). However, as this *in vivo* data came from tissue samples containing tumour and non-cancer cells, it was important to rule out this association as being driven by cancer cell specificity of both methylation and CNV. To assess this, we computed the Spearman correlation coefficients relating each CpG’s methylation β-values to the corresponding tissue sample’s tumour cellularity, and found that these two correlation coefficients were indeed significantly correlated (p<2.2×10^−16^ in both cohorts, Figure 1C-D). Given this finding, we used rank-based partial correlation as a cellularity-adjusted measure of association between methylation and copy number. After adjusting for the potential confounding effects of tumour cellularity, the statistical significance of observed positive partial correlations between DNA methylation and copy number were still dramatically greater than if they were independent for all genes (Figure 1E-F).

**Figure 1.**
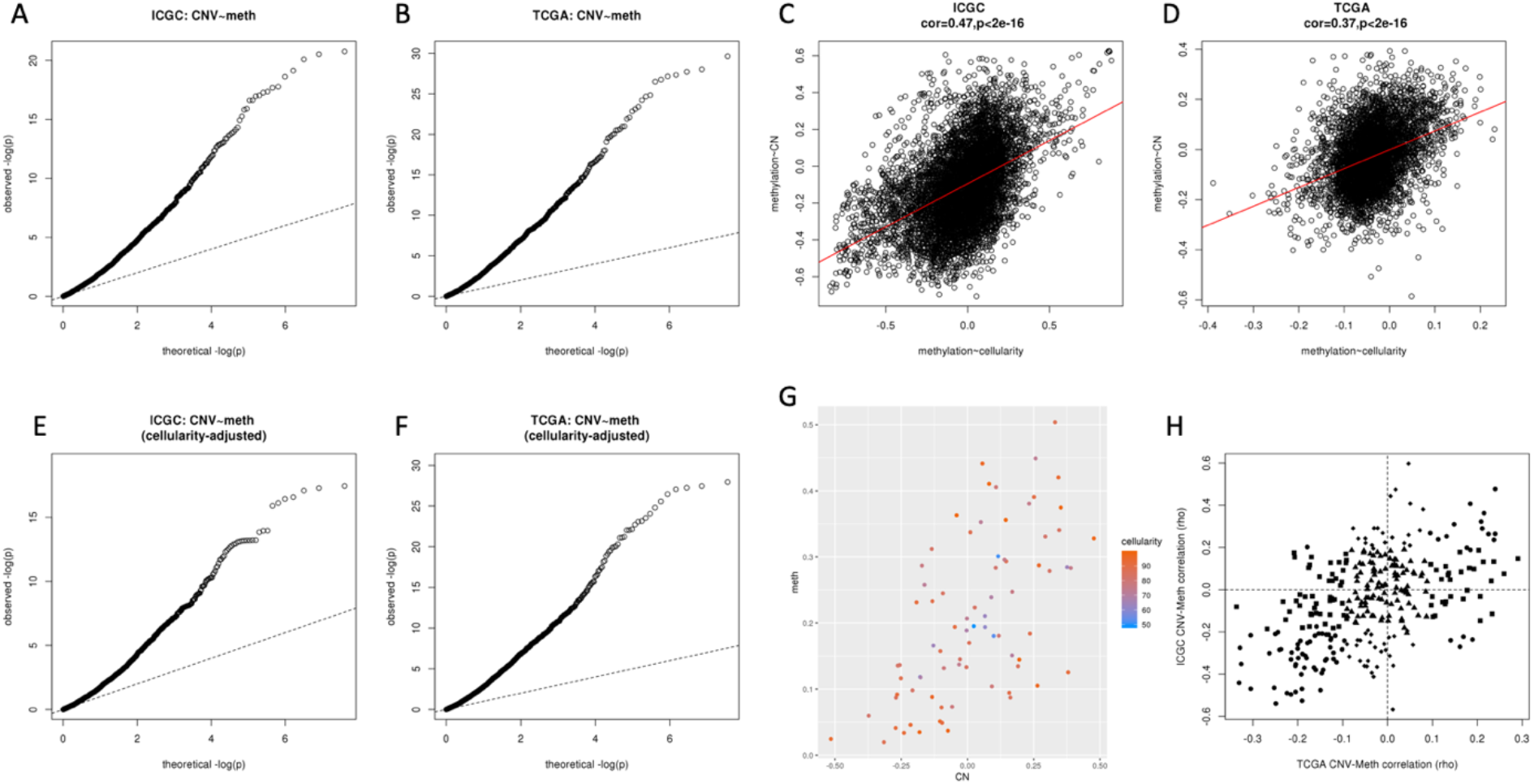
Compensatory DNA methylation in regions of genomic copy-gain. A) Quantile-quantile plot showing the distribution of significance estimates (−log p-values) of positive Spearman correlation coefficients between gCN and DNA methylation in the ICGC dataset, compared to the theoretical distribution under independence of gCN and DNA methylation for all loci (uniformly-distributed p-values, indicated by the dashed line). B) As Fig.1A but for TCGA cohort. C) Plot of correlations between gCN to DNA methylation at each genomic locus against the correlations between DNA methylation and tumour cellularity, ICGC cohort. X-axis shows Spearman correlation coefficient relating gCN estimate to methylation beta-value, y-axis shows Spearman correlation coefficient relating methylation beta-value to cancer cell content of the corresponding tumour sample. Red lines indicate best fit (minimizing sum-squared errors) linear model relating CN~methylation correlations to methylation~cellularity correlations. D) As Fig.1C but for TCGA cohort. E) Quantile-quantile plot showing the distribution of significance estimates (−log p-values) of Spearman partial correlation coefficients between gCN and DNA methylation in the ICGC dataset, controlling for cancer cell content of the corresponding tumour samples. Dashed line gives the theoretical distribution under independence of gCN and DNA methylation, when adjusting for differences in cancer cell content, for all loci (uniformly-distributed p-values). F) As Fig. 1E but for TCGA cohort. G) Plot of DNA methylation beta-value (y-axis) against copy-number (x-axis, as log_2_(CN/2)) for 1 CpG locus (cg06511189) in the FAM58A promoter, for tumour samples in the ICGC cohort. Points are coloured by tumour cellularity (percentage cancer cell content in tissue sample). H) Plot illustrating conservation of locus-wise gCN to DNA methylation correlations across the two HGSOC cohorts analysed. X-axis shows Spearman correlation coefficient relating gCN to methylation in TCGA cohort, y-axis shows Spearman correlation coefficient relating gCN to methylation in ICGC cohort.

To illustrate the correlations between copy-number and methylation (and in some cases tumour cellularity), Figure 1G shows a plot of methylation levels against copy number, coloured by cellularity. Further examples are given in Supplementary Figure S3. A summary of the numbers of loci affected by recurrent gCN gain and compensatory methylation is given in Table 2, and genomic co-ordinates of the individual loci are given in Supplementary Table S1. Each of these analyses reflects the full set of probes lying in promoter regions from the corresponding measurement platform.

**Table 2.** Summary of DCPM events in each cohort

| Cohort | Promoter CpGs Amplified in >10% Tumours | CpGs with copy-number associated methylation | Percentage | CpGs with copy-number associated methylation, excluding cellularity-associated loci | Percentage |
| --- | --- | --- | --- | --- | --- |
| ICGC | 6680 | 635 | 9.5% | 391 | 5.9% |
| TCGA | 5084 | 832 | 16% | 680 | 13% |

The enrichment of correlation between copy-gain and DNA methylation presented in Figure 1 reflects methylation at individual CpG loci. For some CpG islands mapping to regions of recurrent gCN gain, multiple CpG loci are represented in the ICGC methylation dataset. The locus-wise analysis was therefore repeated at the CpG island level: Supplementary Figure S2C shows that an enrichment of correlations between copy-gain and increased average DNA methylation across CpG islands is observed, as at the individual locus level.

We compared the individual correlations for all 303 CpG loci measured in both the two cohorts with sufficient frequency of amplification, observing a similar degree and direction of association between gCN and methylation across the two cohorts (Fig 1H: Pearson correlation coefficient 0.51, p<2×10^−16^). Loci with a significant increase in DNA methylation on increase in gCN (defined as Spearman’s correlation test p<0.1) in one of the cohorts were approximately 3 times as likely to show a statistically significant increase in the other cohort than would be expected by chance (hypergeometric p<1×10^−5^). These observations are important as they suggest that it is a property of these particular genomic loci that result in an increase in methylation upon increase in copy number, as opposed to a general tendency for methylation changes to occur more frequently within regions of genomic copy gain. We refer to the assumed phenomenon giving rise to these associations as Dosage-Compensatory Promoter Methylation (DCPM). It may be noted that the observed negative correlations are also reproducible, therefore certain loci tend to decrease methylation upon increased copy number. As this is likely to have different implications from DCPM, we will focus on DCPM for the remainder of this manuscript.

To confirm our hypothesis that the associations we observed were driven by increase in methylation on genomic copy gain, rather than a decrease in methylation upon copy loss (for instance, methylation of the other allele), the correlation coefficients were recomputed for each locus excluding samples with copy loss. These recomputed copy-neutral vs copy-gain correlations showed similar enrichment and conservation across the two cohorts as those computed across the full cohorts (Supplementary Figure S4A-C). When investigating CpG methylation of loci affected by copy-loss but not copy-gain, we found fewer loci with sufficiently frequent copy number aberration to test, but these were also enriched for positive correlations between gCN and methylation (Supplementary Figure S4D-F). This suggests that regulation of promoter methylation in the event of gCN change could reflect a mechanism cancer cells could harness to prevent amplification-associated over-expression or deletion-associated loss of expression in deleterious passenger genes that are not drivers of the gCN change.

### Dosage-compensatory promoter methylation nullifies expected increase in gene expression upon increase in copy number

Next we investigated the impact of dosage-compensatory DNA methylation on the relationship between genomic copy number and gene expression. Starting with the TCGA cohort, the gene expression measurements for each microarray probe were mapped to their absolute genomic copy number estimate in each sample, and for each probe the Spearman correlation coefficient was calculated to relate the normalized gene expression intensity to copy number estimate across the cohort. The distribution of correlation coefficients across all profiled genes is presented in Fig 2A with the dashed black line, while the corresponding distribution for genes featuring DCPM is indicated with the solid red line. The dashed line shows that most of the genes result in Spearman correlation coefficients greater than zero, implying that increases is gCN generally correspond to increases in gene expression. However, the genes affected by DCPM have less correlation between gCN and expression. Using RNAseq data from the ICGC cohort, a similar approach was taken to evaluate the relationship between gCN and corresponding gene expression across the samples. The distributions of Spearman correlation coefficients are presented in Fig 2B. Again these show that most genes have increases in gCN coinciding with increases in gene expression, but for genes affected by DCPM there is very little association.

**Figure 2.**
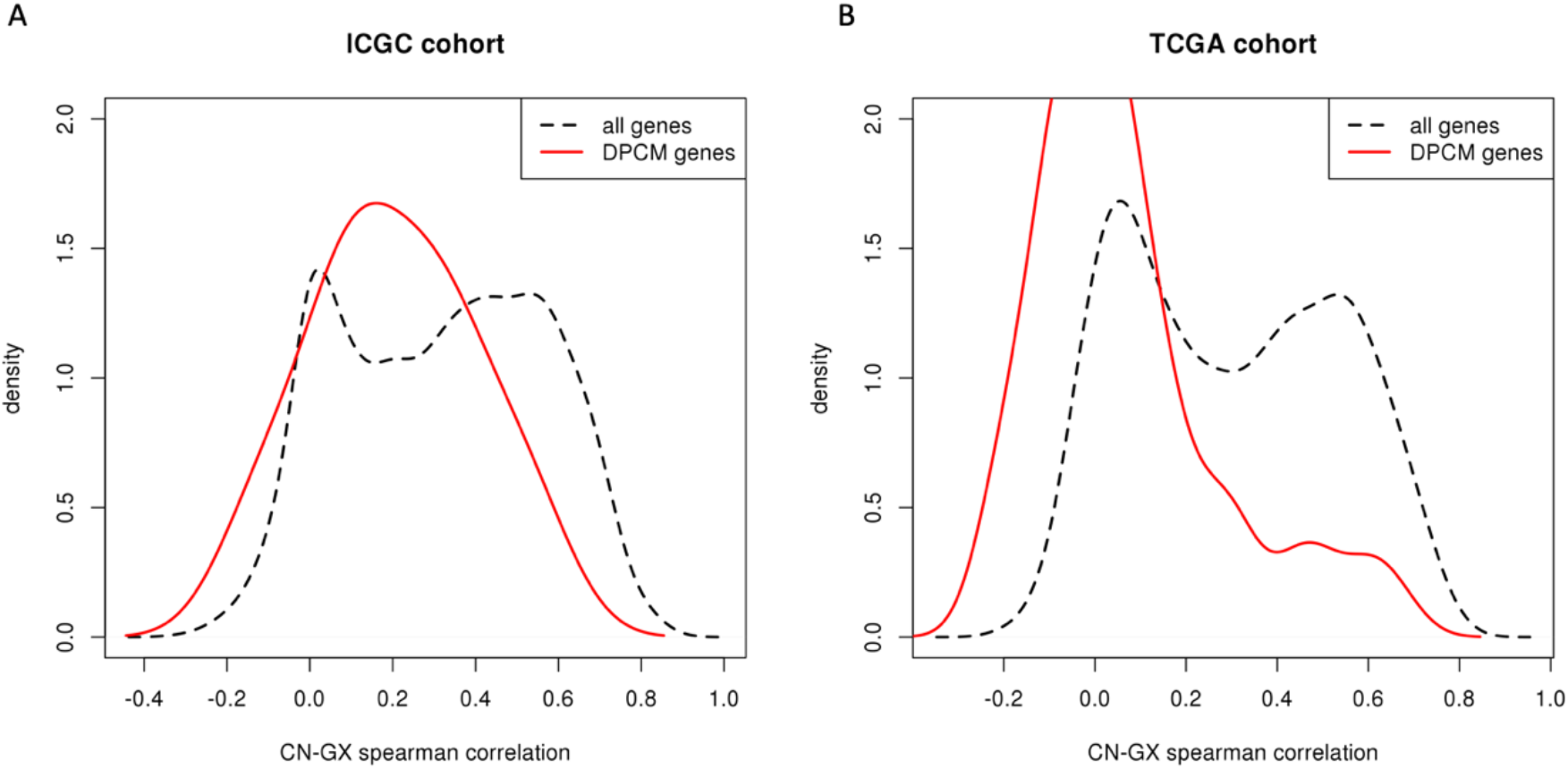
Genes with dose-compensating promoter methylation lose association between copy number and gene expression. A) Loss of association between gCN and gene expression level for genes affected by DCPM, ICGC cohort. Dashed black line gives kernel density estimation for distribution of all gene-wise Spearman correlation coefficients between absolute copy number and normalized expression level. Solid red line gives equivalent distribution of correlation coefficients for the genes affected by DCPM. X-axis shows Spearman correlation coefficient, y-axis shows relative proportion of genes with the corresponding correlation between gCN and gene expression. B) As Fig.2A but for TCGA cohort.

With this evidence we hypothesized that if certain genes exhibit a dose-limiting selective disadvantage to cancer cells, it would encourage increases in promoter methylation whenever these genes were affected by regional genomic copy gain. This could occur when structural variant results in amplification of oncogenic driver genes and confers a sufficient advantage to result in clonal expansion, but expression of linked genes in the amplicon that are negative regulators of tumour cell growth is selected against. A schematic illustrating our proposed explanation for these observations is provided in Fig 3A.

**Figure 3.**
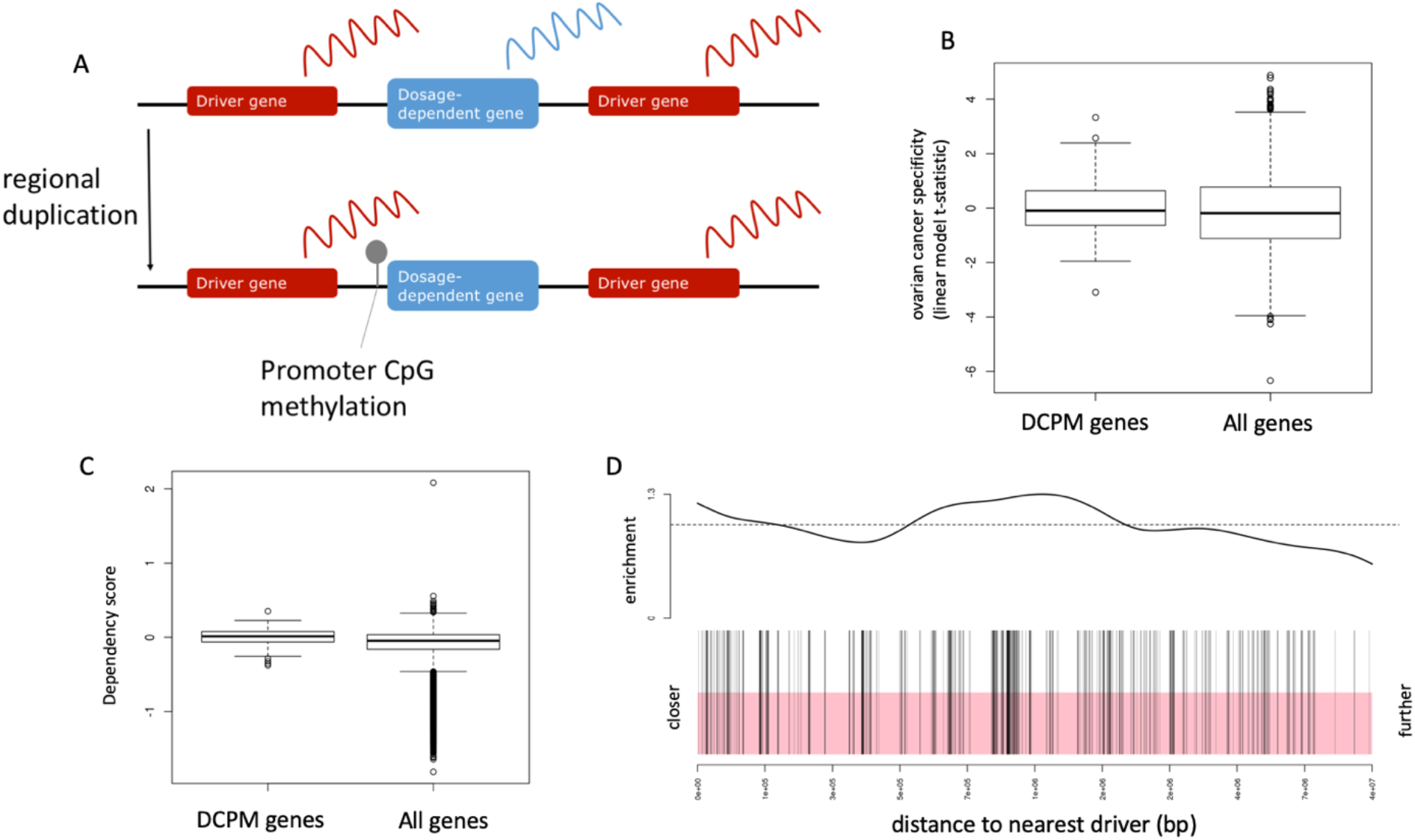
Dosage-compensatory promoter methylation as a mechanism to overcome selective disadvantage introduced by increasing copy number of dosage-dependent passenger genes. A) Schematic illustrating hypothesis for DCPM as a means of counteracting negative selective pressure from genes with dosage-limiting effects located within amplicon containing oncogenic driver genes. Line represents a genomic region with a dosage-dependent gene (blue) flanked by two oncogenic driver genes (red). When the region is duplicated, an advantage is gained from increased expression of the oncogenic drivers, but a disadvantage arises from increased expression of the dosage-dependent gene. Combined, this encourages selection of cells that have increased the genomic copy number in this region but acquire CpG methylation in the promoter of the dosage-dependent gene. B) Ovarian cancer cell line specificity of selective pressure on shRNA incorporation from ACHILLES screen. Box plots show distribution of t-statistics for increase in selective pressure for incorporating each of a library of shRNAs into ovarian cancer cell lines, compared to other cancer cell lines. The distribution for shRNAs targeting genes affected by DCPM is given by the left-hand box, right-hand box gives distribution for all shRNAs targeting any gene. The shift towards higher values of t-statistics in the left-hand box implies that incorporation of shRNAs targeting genes affected by DCPM tends to confer a greater selective advantage in ovarian cancer cell lines when compared to other cancer cell lines. Box plot elements defined as follows: center line, median; box limits, upper and lower quartiles; whiskers, 1.5x interquartile range; points, outliers). C) Selective pressure on CRISPR knock-outs targeting DCPM genes, from DepMap cancer cell dependency screen. Box plots show distribution of median CERES dependency scores for incorporating CRISPR sgRNAs targeting each gene into all profiled cancer cell lines. The distribution for sgRNAs targeting genes affected by DCPM is given by the left-hand box, right-hand box gives distribution for all sgRNAs targeting any gene. The higher values of CERES scores in the left-hand box implies that incorporation of sgRNAs targeting genes affected by DCPM tends to confer a selective advantage relative to incorporating sgRNAs targeting genes not affected by DCPM. Box plot elements defined as follows: center line, median; box limits, upper and lower quartiles; whiskers, 1.5x interquartile range; points, outliers). D) Barcode plot showing tendency of DCPM genes to be closer to their nearest COSMIC driver gene than would be expected for an equivalent set of randomly-chosen genes. Bottom panel shows distance to nearest driver gene for all genes, ranked from closest (left) to furthest (right). Vertical black lines indicate rank of DCPM genes. Top panel shows enrichment as ratio of observed number of DCPM genes ranked above each position, relative to number expected from random sampling. Horizontal dashed line shows observed/expected ratio = 1 (i.e. no enrichment). Genes affected by DCPM are systematically closer to COSMIC census genes than expected by chance (mean-rank gene set enrichment test p<10^−4^).

### Silencing of genes affected by dosage-compensatory promoter methylation appears to confer a selective advantage specific to ovarian cancer cell lines

If overexpression of these candidate genes were to confer a selective disadvantage to tumour development, it might be expected that cells harbouring extra copies of these genes would acquire a proliferative advantage when incorporating shRNAs or CRISPR sgRNAs targeting those same genes. The Achilles project (Cowley et al. 2014) quantified incorporation of pooled shRNA libraries in 216 cancer cell lines following 72 hours of subsequent growth in media, giving estimates for the relative vulnerability or selectivity of each cell line to knockdown of each gene. We compared the relative incorporation of shRNAs into ovarian cancer cell lines, compared to all other cell lines. Interestingly, the set of genes with substantial DCPM are significantly biased towards high shRNA incorporation scores in cell lines with ovarian origin (Wilcoxon rank sum p<0.028) as shown in Fig 3B. This represents a systematic enrichment of outgrowth of clones that incorporate shRNAs targeting these genes, specifically in ovarian cancer cell lines. The DepMap project (Meyers et al. 2017) quantified incorporation of pooled sgRNA libraries in 324 cancer cell lines, giving estimates of the relative vulnerability or selectivity of each cell line to knockout of each gene. The set of genes with substantial DCPM are significantly biased towards high median sgRNA incorporation scores across the whole DepMap dataset (Wilcoxon rank sum p<1×10^−6^) as shown in Fig 3C.

Our hypothesis involves proximity to oncogenic drivers as the underlying cause of recurrent copy gain affecting these genes, providing a stronger selective pressure than the apparent disadvantage which can be overcome by epigenetic silencing of the affected genes. To test this aspect of the hypothesis we computed the distances from the DCPM candidates (discovered across the two cohorts of primary tumour samples) to the nearest COSMIC census cancer gene (Forbes et al. 2011). Among all genes tested for DCPM, the set affected by DCPM were systematically closer to COSMIC census genes (mean-rank gene set enrichment test(Michaud et al. 2008) p<10^−4^, Figure 3D).

### Forced over-expression of genes affected by dosage-compensatory methylation decreases tumour spheroid formation efficiency of ovarian cancer cell lines

If compensatory methylation is employed by cancer cells to reduce a selective disadvantage of overexpressing certain genes, one would expect that exogenous overexpression would impair phenotypic traits of cancer cells that relate to tumour growth *in vivo*. To test this hypothesis, we selected candidate genes in the 3q12 locus, which has copy number gain in the high-grade serous ovarian cancer cell lines PEA1 and PEO1 (Figures 4A&5A). The candidate genes GPR171 & ZMAT3 were selected because in both patient cohorts they were frequently amplified, did not show significantly increased gene expression on copy-gain, featured at least 1 CpG locus mapping to the gene with significantly increased methylation on copy-gain, and were flanked by 2 known oncogenic drivers each (SIAH2, RAP2B, ECT2 & PIK3CA shown in Figures 4B&5B). Baseline gene expression for the candidates GPR171 & ZMAT3, and for the oncogenic drivers SIAH2, RAP2B, ECT2 & PIK3CA were measured by qRT-PCR in the ovarian cancer cell lines and compared to the non-malignant fallopian tube derived line FT190 (Supplementary Table S2). FT190 cell line was used as HGSOC is thought predominantly to arise from fallopian tube(Labidi-Galy et al. 2017). It can be seen that both candidate genes were detectable in all cell lines assayed, although GPR171 is expressed at quite a low level. Figure 4C shows increased expression of ECT2 and decreased expression of both GPR171 and ZMAT3 in PEA1 (relative to FT190). Figure 5C shows increased expression of all 4 oncogenic drivers and decreased expression of ZMAT3 in PEO1 (relative to FT190). We used bisulphite pyrosequencing to assay methylation in these cell lines (Supplementary Table S3), finding only one CpG site for ZMAT3, and none for GPR171, that showed increased methylation in the HGSOC cell lines relative to FT190 (Figures 4D&5D). The relative repression of these genes in the 2 profiled HGSOC cell lines might arise from a mechanism other than DNA methylation, but they still serve as suitable models to evaluate the selective advantage of preventing over-expression of DCPM genes affected by copy-gain. Sub-clones of PEA1 and PEO1 cell lines were created that stably over-express each of the candidate DCPM genes, with clear increases in corresponding mRNA levels (Figures 4E&5E).

**Figure 4.**
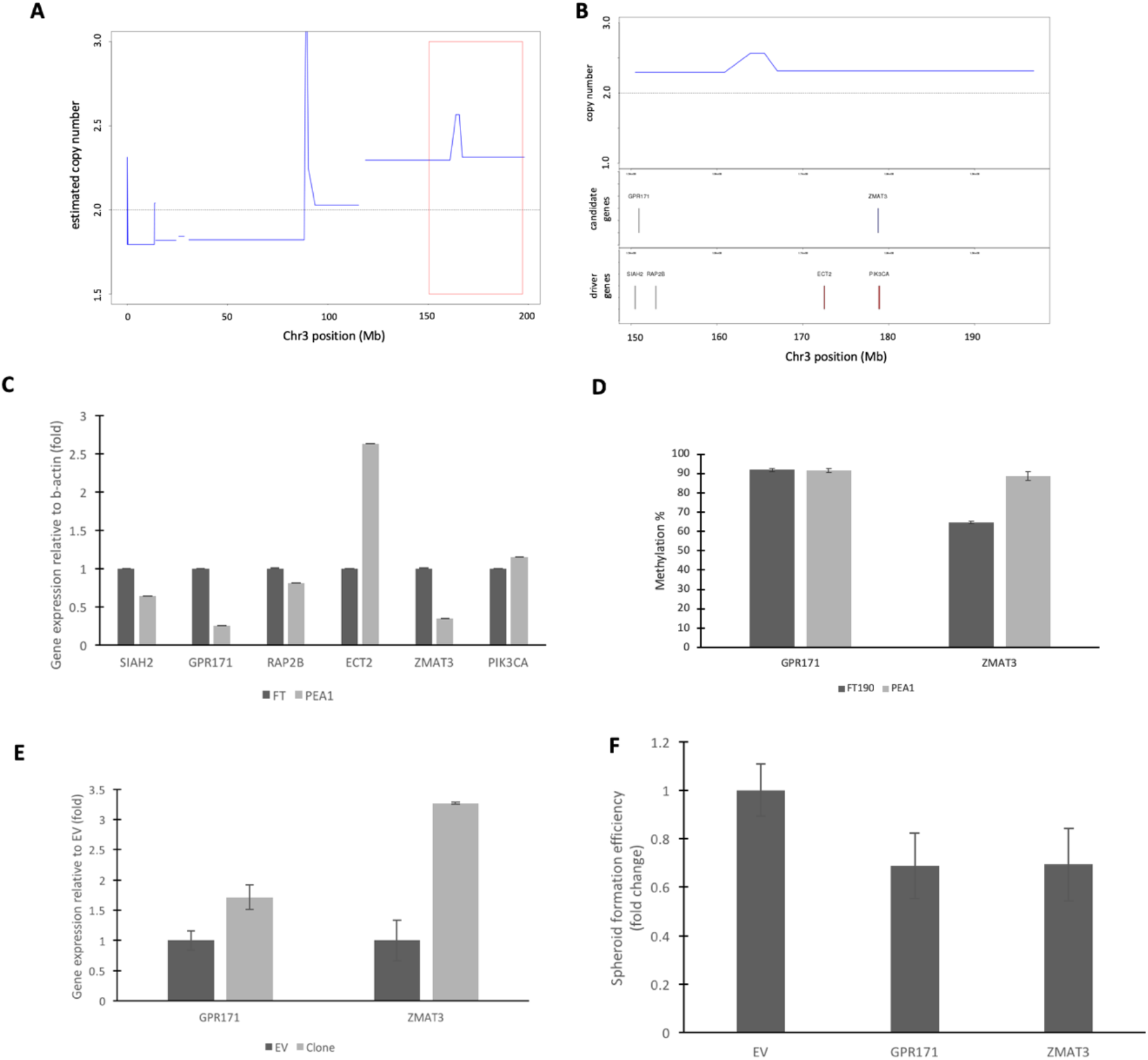
Characterizing consequences of DCPM candidate overexpression in PEA1. A) Chromosome 3 copy-number profile for PEA1 cell line, showing SNP-array derived segmented copy-number estimate at each position. B) Copy-number profile zooming in on highlighted region of large 3q amplicon featuring candidate loci with compensatory methylation and potential driver oncogenes. C) Baseline gene expression of candidate genes and drivers in ovarian cancer cell line PEA1 relative to fallopian tube derived FT190. Values shown represent fold change relative to FT190 values of qRT-PCR measurement for each gene normalized to β-actin. Data shown is a mean of 3 biological repeats. Standard error bars shown. D) CpG methylation of candidate genes in ovarian cancer cell line PEA1 and fallopian tube derived FT190. Values shown represent mean methylation percentage of 3 independent repeats. Error bars give standard deviation across repeats. E) Gene expression of PEA1-derived clones over-expressing candidate genes. Values shown represent qRT-PCR measurements normalized to β-actin, relative to PEA1 cell line transfected with an empty vector. Data shown is a mean of 3 technical repeats. Standard error bars shown. F) Spheroid formation efficiency evaluated from a spheroid count (>50μm) from 4 replication cycles (4 days) of the same density of cells whilst grown in full medium under optimal conditions in an ultra-low adhesion 6-well dish. Y-axis gives fold change from the empty vector (EV). Data shown is a mean of 4 independent experiments (standard error bars shown), each constituting 3 technical repeats.

**Figure 5.**
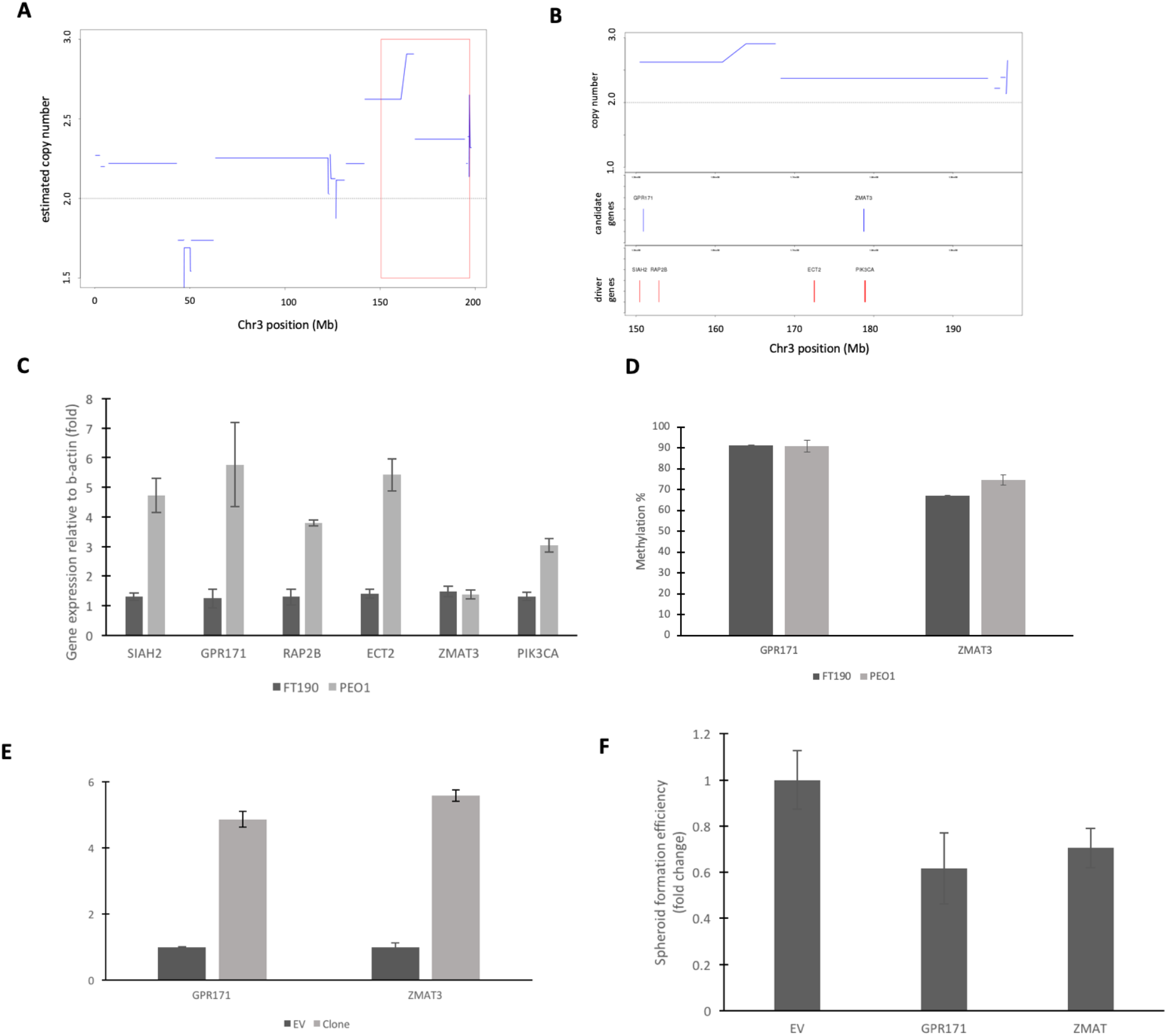
Characterizing consequences of DCPM candidate overexpression in PEO1. A) Chromosome 3 copy-number profile for PEO1 cell line, showing SNP-array derived segmented copy-number estimate at each position. B) Copy-number profile zooming in on highlighted region of large 3q amplicon featuring candidate loci with compensatory methylation and potential driver oncogenes. C) Baseline gene expression of candidate genes and drivers in ovarian cancer cell line PEO1 relative to fallopian tube derived FT190. Values shown represent fold change relative to FT190 values of qRT-PCR measurement for each gene normalized to β-actin. Data shown is a mean of 6 biological repeats. Standard error bars shown. D) CpG methylation of candidate genes in ovarian cancer cell line PEO1 and fallopian tube derived FT190. Values shown represent mean methylation percentage of 3 independent repeats. Error bars give standard deviation across repeats. E) Gene expression of PEO1-derived clones over-expressing candidate genes. Values shown represent qRT-PCR measurements normalized to β-actin, relative to PEO1 cell line transfected with an empty vector. Data shown is a mean of 3 technical repeats. Standard error bars shown. F) Spheroid formation efficiency evaluated from a spheroid count (>50μm) from 4 replication cycles (4 days) of the same density of cells whilst grown in full medium under optimal conditions in an ultra-low adhesion 6-well dish. Y-axis gives fold change from the empty vector (EV). Data shown is a mean of 3 independent experiments (standard error bars shown), each constituting 3 technical repeats.

Figure 4F shows that for both candidate genes, overexpression resulted in a significant decrease in spheroid formation efficiency in PEA1 (t-test p<0.05). Figure 5F shows similar results for PEO1. Anchorage-independent growth of cells in spheroids is generally taken as a measure of increased cell transformation. Effects on this phenotype by DPCM gene overexpression provides clear evidence that overexpression of these genes would result in a selective disadvantage for tumour growth and development, supporting a model in which cancer cells experience pressure to acquire aberrant promoter hypermethylation at dosage-sensitive genes in genomic proximity to oncogenic drivers.

## DISCUSSION

Our genome-wide integrative analysis of genomic copy number, DNA methylation and gene expression data led to discovery of dosage-compensating promoter methylation for a subset of genes. This finding was replicated across two independent cohorts of HGSOC. We found some loci where the association appears to be driven by amplification of genes with tumour-specific methylation. Some previous studies have reported associations between copy number and DNA methylation in tumour tissues, at genes where copy number is not associated to gene expression (Flanagan et al. 2010; Murria et al. 2015; Mohanty et al. 2017). Our findings suggest that genes regulated by DNA methylation lying within amplicons in tissue samples with variable tumour cellularity could bias these previous studies, and it is therefore important to adjust for cellularity-driven effects. The fact that the correlation coefficients for each locus are significantly correlated across the two cohorts suggests that DCPM is a feature of specific loci, rather than a phenomenon that occurs repeatably but randomly throughout the genome. We have restricted our analysis to loci in promoter CpG islands, as the functional link to silencing of gene expression is well understood (Bird and Wolffe 1999). Our analysis has also been limited to the specific CpG sites measured on the Illumina HumanMethylation27k and 450k microarray platforms. We have established that these CpG sites are loosely representative of other CpG sites within their respective CpG islands, but it would be interesting to evaluate more comprehensively the consistency of DCPM across many CpG sites within each promoter CpG island affected by DCPM. There is evidence that not all promoter CpG islands are affected by methylation in the same way (Weber et al. 2007). This implies that a more comprehensive characterization of DCPM could focus on better coverage of CpG sites across certain classes of promoter, notably those with *de novo* methylation associating with loss of expression. Furthermore, a recent systematic analysis of transcription factor DNA-binding has shown methylation-specific binding of certain transcription factors to certain sequences of DNA (Yin et al. 2017). If whole-genome CpG methylation data were available, it might be possible to identify systematic dosage-compensatory motif methylation (or de-methylation) for particular transcription factors, linking non-promoter CpG methylation to expected functional consequences in cancer cells. At a genome-wide level, overall increases in ploidy would increase the copy-number of any dose-limiting genes. It would therefore be interesting to check if this coincided with a global increase in promoter methylation, where separate quantification of ploidy for tumour sample is available. We have not investigated the potential allele-specificity of DCPM events, as this would require sequence-specific genome-wide analysis of methylation through whole-genome bisulphite sequencing. Evaluating whether or not DCPM events tend to occur on the allele affected by copy-gain could give some insight into potential mechanisms of observed DCPM.

We had initially hypothesized that genes might be held in dosage-compensating regulation if their protein products were involved in reactions sensitive to stoichiometric ratios. It has been suggested that most metabolic pathways and developmental processes are gene dosage-dependent (Birchler et al. 2001), but we did not see a particular enrichment for genes annotated with metabolic functions in the DCPM targets. In fact, we didn’t find enrichment of any particular biological processes in the DCPM-affected genes, which may indicate this as a broadly-applicable mechanism affecting genes across any biological processes that cancer cells manipulate to proliferate in their environment. The fact that the DCPM-affected genes were enriched for ovarian cancer specific selection of incorporated shRNAs in the Achilles project, and were generally enriched for sgRNA incorporation in the DepMap project, is particularly reassuring in support of the hypothesis that DCPM indicates a gene that is disadvantageous when overexpressed. The fact that different loci will be affected in different patients, due to differing genomic copy-number landscapes, could suggest a potential mechanism for nonspecific epigenetic therapies treating cancers by disrupting DNA methylation genome-wide. By this logic, reversal of the DCPM effect through epigenetic therapies might be expected to impair growth of cancer cells featuring extensive genomic copy gains.

In case the acquisition of methylation at individual amplified promoters impacts on patient prognosis, we tested the association of methylation with both progression-free and overall survival, amongst patients harbouring an amplification. But quantile-plots of the p-values from the resulting regression models revealed no noticeable deviation from the expected distribution under randomness (Supplementary Figure S5). This should not really be surprising, as patient outcomes will depend on response to treatments they will be exposed to after the methylation data was collected. These treatments will impose different selection pressures and may result in DCPM at different loci.

We also found marked enrichment for loci showing negative correlation between copy-number and methylation. We first considered this an obvious consequence of driver genes promoting tumour formation being over-expressed through both copy gain and loss of methylation. However, it may be interesting to consider an implication that there is a greater selection pressure for loss of methylation when the driver gene is amplified: perhaps the amplification results in (or is a result of) addiction to over-expression of the gene in question?

Why do the sets of genes affected by DCPM (which are a minority among genes with recurrent copy gains) show consistency across two independent cohorts? This suggests that there is a general selection pressure for certain genes to be affected by copy gain, and additional selection for preventing over-expression of a subset of these genes. It is of note that these consistently-affected loci showed systematic bias to preferential selection for knock-down specifically in ovarian cancer cell lines. As the affected genes were found to be genomically closer to known driver oncogenes than would be expected by chance, it is likely that the consistency of DCPM loci arises from an initial selection for cells harbouring increased copy number of a nearby driver gene, and then subsequent selection for cells that acquire epigenetic silencing of a particularly dosage-sensitive over-expressed passenger gene. We have demonstrated that forced over-expression of two genes affected by compensatory methylation in HGSOC results in a decrease in spheroid formation efficiency. This indicates a specific selective advantage conferred to cells that prevent copy gain associated over-expression of these genes. With these observations in mind, we speculate that the fundamental molecular pathology of HGSOC might be influencing which genes have such a dosage-specific effect that DCPM becomes selected for in a locus-specific manner. It would be interesting to extend our approach to other cancer types, and perhaps even to restrict analyses to within relatively homogeneous molecular subtypes, in order to compare disease-specific DCPM selectivity. Such analysis could shed light on the aspects of equilibrium maintained in cancers despite loss of regulatory pathways and checkpoints, and reveal vulnerabilities of cancers with a wide range of molecular pathologies underpinned by variations in genomic copy-number.

In conclusion, we have identified dosage-compensatory promoter methylation (DCPM) as a recurrent epigenetic phenomenon enabling ovarian cancer cells to avoid overexpression of certain genes that have increased genomic copy number, possibly arising as a passenger effect of proximity to driver oncogenes. We have demonstrated that overexpressing such genes in two ovarian cancer cell lines harbouring increased copy number of those loci decreases tumour spheroid formation. This supports our hypothesis that dosage-compensatory methylation may be invoked to promote cancer growth.

## METHODS

### Datasets

For the TCGA cohort, genome-wide copy-number profiles were obtained from Affymetrix SNP6 microarrays processed to segmented copy-number estimates with circular binary segmentation (Olshen et al. 2004) as implemented in the R package *DNAcopy* available from Bioconductor(Gentleman and \it e1 al 2004). DNA methylation profiles were obtained from Illumina HumanMethylation27k microarrays ready preprocessed (Level 2) as probe-wise methylation beta values. Gene expression profiles were obtained from Affymetrix HT-HGU133A microarrays ready preprocessed (Level 2) as probe-wide log2 intensity values. Tumour cellularity of each tissue sample was computed as median cellularity of each slide from the corresponding sample, taken from the annotation file ‘clinical_slide_all_ov.txt’. All these datasets were downloaded from the TCGA data portal, and are available from the legacy archive at the NCI’s Genomic Data Commons: https://portal.gdc.cancer.gov/legacy-archive For the ICGC cohort, genome-wide copy-number profiles were obtained from Affymetrix SNP6 microarrays. DNA methylation profiles were obtained from Illumina HumanMethylation450k microarrays. Gene expression profiles were obtained from RNA-seq libraries. All data was processed as described in (Patch et al. 2015a). Tumour cellularity of each tissue sample was obtained from sample annotation file ‘sample.OV-AU.tsv’. All these datasets are available from the ICGC data portal https://dcc.icgc.org with project title ‘OV-AU’.

Project Achilles data were downloaded from https://portals.broadinstitute.org/achilles as a matrix of ATARIS-processed shRNA incorporation scores, as described (Cowley et al. 2014). DepMap data were downloaded from https://depmap.org/portal/download/ as a matrix of batch corrected CERES gene effect scores, as described (Meyers et al. 2017).

Genome-wide copy-number profile from PEA1 and PEO1 cell lines were obtained from Illumina Human1M-Duov3 DNA Analysis BeadChip microarray data downloaded from Gene Expression Omnibus with accession GSM459851 and GSM459853, respectively. Downloaded log ratio values were transferred from into hg19 (GRCh37.p13) co-ordinates using UCSC liftOver tool implemented in *rtracklayer* R package in Bioconductor. Probe-level copy-number log-ratio values were smoothed and segmented using the *DNAcopy* package and then mapped to genes using the Ensembl hg19 (GRCh37.p13) annotations.

### Statistical Analysis

DNA methylation β-values (proportion of DNA molecules with the corresponding CpG methylated) were analysed for individual CpG loci, each located within promoter CpG islands. Spearman rank correlation coefficients were calculated to relate DNA methylation β-values to absolute copy number estimates, with statistical significance of any observed correlation evaluated using the *cor.test* function in **R**. Enrichment over expected correlation was calculated assuming a uniform distribution of p-values under the null hypothesis of no correlations between methylation and copy number. Partial correlations were evaluated for each locus using linear regression models fitting the rank of DNA methylation β-values in each tumour sample as a function of the rank of tumour cellularity (cancer cell content of the tissue sample) and rank of genomic copy-number at the locus. Coefficients and p-values were obtained from linear models through the *summary.lm* function in **R**.

Multiple hypothesis testing adjustment was performed using the Benjamini-Hochberg method implemented in the *p.adjust* function in **R**. Overlap between DCPM loci identified from the two cohorts was evaluated using a hypergeometric test, based on the distribution implemented in **R**. To establish the extent to which methylation levels of individual CpG loci are representative of the CpG islands, for each CpG island we computed the median Pearson Correlation Coefficient between each pair of CpG loci within the island. The distribution of these median intra-island correlation coefficients is given in Supplementary Figure S6, showing that the methylation levels of the majority of loci are positively correlated with the levels of other loci in the same CpG island. Systematic shift in preferential shRNA incorporation from the ovarian cancer cell lines in the Achilles dataset was tested by first evaluating empirical Bayes moderated t-statistics comparing each gene’s ATARIS-preprocessed shRNA incorporation scores in the ovarian cancer cell lines against the gene’s equivalent shRNA incorporation scores across all non-ovarian cancer cell lines. The resulting t-statistics were used as the basis for a Wilcoxon rank sum test of systematic bias towards high ranking of the DCPM loci, implemented using the *geneSetTest* function within the **R** *limma* package, distributed in Bioconductor.

Systematic shift in preferential incorporation of sgRNAs targeting DCPM affected genes from the DepMap dataset was evaluated using Wilcoxon rank sum test on the CERES scores, using all non DCPM genes for comparison.

### Cell Lines

Cell lines were obtained from the cell stocks of the Department of Cancer and Surgery. All cell lines were *Mycoplasma-free*. Fallopian tube epithelial secretory cell line FT190 was obtained from Dr Paula Cunnea, gifted from Alison Kast and Ronny Drapkin (Karst et al. 2011). PEA1 is an adherent mammalian cell line derived from a high grade serous ovarian cancer patient prior to cisplatin (chemotherapy) treatment. PEO1 is an adherent mammalian cell line derived from a high grade serous ovarian cancer patient following chemotherapy (cisplatin, 5-Fluorouracil and chlorambucil) treatment but prior to development of clinical resistance to the chemotherapy agents (Langdon et al. 1988). PEA1 cell line was kept at 37°C, 5% CO_2_, in RPMI-1640 medium (Sigma-Aldrich, St.Louis, US) supplemented with 2mM L-glutamine (Gibco) and 10% FCS and 100 units/mL Penicilium and 100 μg/mL Streptomycin (Gibco). To generate overexpressing cells, each cell line (PEA1 and PEO1) was transfected using Fugene^®^ with the following overexpressing plasmids: *ZMAT3* (RC202508, Origene), *GPR171* (RC207757, Origene) or the empty vector control (EV)(PS100001, Origene). PEA1 clones were kept under identical conditions and supplemented medium replacing PenStrep with 500μg/ml Geneticin^®^ (Life technologies).

### Quantitative Real-Time PCR (qRT-PCR) for gene expression

qRT-PCR was performed in order to obtain relative gene expression values. RNA used was using QIAGEN^®^ RNeasy Mini Kit and cDNA synthesised utilising the High Capacity cDNA Reverse Transcription Kit (Applied Biosystems). 10ng of cDNA was used with the designed primers (Supplementary Table S4) to quantify the desired gene expression. Values obtained analysed using the ΔΔCt method, normalised to β-actin.

### Bisulphite Pyrosequencing

A total amount of 500 ng of genomic DNA was bisulphite modified using the Zymo Research EZ-DNA Methylation Kit (Cambridge Bioscience) according to the manufacturer’s instructions. Sequences of pyrosequencing primer sets are provided in Supplementary Table S4. Pyrosequencing PCR was performed in duplicate for each sample in a 25 μl volume containing an end-concentration of 1 U Faststart Taq polymerase (Roche, Welwyn Garden City, UK), 1x FastStart PCR Buffer including 2 μM MgCl_2_ (Roche), 0.05 mm dNTPs (Roche), 0.4 μM primers (each) adding 1 μl of modified DNA template using the following conditions: 95 °C for 6 min, 45 cycles of 95 °C for 30 s, 58 °C for 30 s, 72 °C for 30 s, followed by 72 °C for 5 min and terminating at 4°C. Pyrosequencing of PCR products was performed on PyroMark Q 96 MD using the PyroGold Reagent Kit (Qiagen) according to the manufacturer’s instructions. The methylation percentage of CpG sites for individual genes was calculated by using the Pyro Q-CpG software (version 1.0.9), Biotage (Uppsala, Sweden).

### Tumour-spheroid formation assay

Tumour-spheroid formation assay was performed using a protocol from (Rizzo et al. 2011). Cells were seeded at an optimised density of 2000 cells/well of a 6-well Ultra low-adhesion plate and kept in the incubator at optimal conditions for 4 replication cycles (4 days). Tumour spheroids were counted by manual inspection of each entire well using Brightfield microscopy. Only spheroids with diameter larger than 50μm were counted. Spheroid formation capacity was calculated by divided the number of spheroids formed by the number of cell plated (2000 cells).

## DATA ACCESS

Genome-wide profiles of copy-number, DNA methylation and gene expression that support the findings of this study are available in the public domain, with instructions for access given in the ‘*Datasets*’ paragraph in the Methods section of this manuscript. For convenience, all processed datasets are made available through Mendeley Data at http://dx.doi.org/10.17632/623s8b9vb4.1 Experimental data presented in this study are available from the corresponding author upon reasonable request. R code used to perform all analyses presented in this study are publicly available in the GitHub repository: https://github.com/edcurry/CNV-compensatory-methylation

## ACKNOWLEDGEMENTS

This study was supported by a Cancer Research UK grant to RB (A13086), Ovarian Cancer Action Research Centre, Imperial Experimental Cancer Medicine Centre, and the Imperial NIHR Biomedical Research Centre. We would like to acknowledge support of Prof David Bowtell and the ICGC Ovarian Cancer study group for access to data from clinical samples and discussion of the work in its initial stages, and Dr Paula Cunnea for provision of FT190 cell line.

## REFERENCES

Akavia UD, Litvin O, Kim J, Sanchez-Garcia F, Kotliar D, Causton HC, Pochanard P, Mozes E, Garraway LA, Pe’er D. 2010. An integrated approach to uncover drivers of cancer. Cell 143:1005–1017.

Baylin SB, Jones PA. 2011. A decade of exploring the cancer epigenome — biological and translational implications. Nat Rev Cancer 11: 726–734.

Bergamaschi A, Kim YH, Wang P, Sørlie T, Hernandez-Boussard T, Lonning PE, Tibshirani R, Børresen-Dale AL, Pollack JR. 2006. Distinct patterns of DNA copy number alteration are associated with different clinicopathological features and gene-expression subtypes of breast cancer. Genes, Chromosomes and Cancer 45:1033–1040.

Beroukhim R, Mermel CH, Porter D, Wei G, Raychaudhuri S, Donovan J, Barretina J, Boehm JS, Dobson J, Urashima M. 2010. The landscape of somatic copy-number alteration across human cancers. Nature 463:899–905.

Birchler JA, Bhadra U, Bhadra MP, Auger DL. 2001. Dosage-dependent gene regulation in multicellular eukaryotes: implications for dosage compensation, aneuploid syndromes, and quantitative traits. Developmental biology 234:275–288.

Bird AP, Wolffe AP. 1999. Methylation-induced repression—belts, braces, and chromatin. Cell 99:451–454.

Bonito NA, Borely J, Wilhelm-Benartzi C, Ghaem-Maghami S, Brown R. 2016. Epigenetic regulation of the homeobox gene MSX1 associates with platinum resistant disease in high grade serous epithelial ovarian cancer. Clinical Cancer Research: clincanres. 1669.2015.

Chang AY-F, Liao B-Y. 2011. DNA Methylation Rebalances Gene Dosage after Mammalian Gene Duplications. Molecular Biology and Evolution 29:133–144.

Ciriello G, Miller ML, Aksoy BA, Senbabaoglu Y, Schultz N, Sander C. 2013. Emerging landscape of oncogenic signatures across human cancers. Nature genetics 45:1127–1133.

Consortium T. 2011a. Integrated genomic analyses of ovarian carcinoma. Nature 474:609–615.

Consortium TCGA. 2011b. Integrated genomic analyses of ovarian carcinoma. Nature 474:609–615.

Cowley GS, Weir BA, Vazquez F, Tamayo P, Scott JA, Rusin S, East-Seletsky A, Ali LD, Gerath WF, Pantel SE. 2014. Parallel genome-scale loss of function screens in 216 cancer cell lines for the identification of context-specific genetic dependencies. Scientific data 1.

Flanagan JM, Cocciardi S, Waddell N, Johnstone CN, Marsh A, Henderson S, Simpson P, da Silva L, Khanna K, Lakhani S. 2010. DNA methylome of familial breast cancer identifies distinct profiles defined by mutation status. The American Journal of Human Genetics 86:420–433.

Forbes SA, Bindal N, Bamford S, Cole C, Kok CY, Beare D, Jia M, Shepherd R, Leung K, Menzies A et al. 2011. COSMIC: mining complete cancer genomes in the Catalogue of Somatic Mutations in Cancer. Nucleic Acids Research 39:D945–D950.

Gentleman RC, \it e1 al. 2004. Bioconductor: open software development of computational biology and bioinformatics. Genome Biology 5:R80.

Karst AM, Levanon K, Drapkin R. 2011. Modeling high-grade serous ovarian carcinogenesis from the fallopian tube. Proceedings of the National Academy of Sciences 108:7547–7552.

Labidi-Galy SI, Papp E, Hallberg D, Niknafs N, Adleff V, Noe M, Bhattacharya R, Novak M, Jones S, Phallen J. 2017. High grade serous ovarian carcinomas originate in the fallopian tube. Nature communications 8:1093.

Langdon SP, Lawrie SS, Hay FG, Hawkes MM, McDonald A, Hayward IP, \it e1 al. 1988. Characterization and properties of nine human ovarian adenocarcinoma cell lines. Cancer Res 48:6166–6172.

Lengyel E. 2010. Ovarian cancer development and metastasis. The American journal of pathology 177:1053–1064.

McCarroll SA, Hadnott TN, Perry GH, Sabeti PC, Zody MC, Barrett JC, Dallaire S, Gabriel SB, Lee C, Daly MJ. 2006. Common deletion polymorphisms in the human genome. Nature genetics 38:86–92.

Meyers RM, Bryan JG, McFarland JM, Weir BA, Sizemore AE, Xu H, Dharia NV, Montgomery PG, Cowley GS, Pantel S. 2017. Computational correction of copy number effect improves specificity of CRISPR–Cas9 essentiality screens in cancer cells. Nature genetics 49:1779.

Michaud J, Simpson KM, Escher R, Buchet-Poyau K, Beissbarth T, Carmichael C, Ritchie ME, Schütz F, Cannon P, Liu M. 2008. Integrative analysis of RUNX1 downstream pathways and target genes. BMC genomics 9:363.

Mohandas T, Sparkes R, Shapiro L. 1981. Reactivation of an inactive human X chromosome: evidence for X inactivation by DNA methylation. Science 211:393–396.

Mohanty V, Akmamedova O, Komurov K. 2017. Selective DNA methylation in cancers controls collateral damage induced by large structural variations. Oncotarget 8:71385.

Murria R, Palanca S, de Juan I, Egoavil C, Alenda C, García-Casado Z, Juan MJ, Sánchez AB, Santaballa A, Chirivella I. 2015. Methylation of tumor suppressor genes is related with copy number aberrations in breast cancer. Am J Cancer Res 5:375–385.

Olshen AB, Venkatraman E, Lucito R, Wigler M. 2004. Circular binary segmentation for the analysis of array-based DNA copy number data. Biostatistics 5:557–572.

Patch A-M, Christie EL, Etemadmoghadam D, Garsed DW, George J, Fereday S, Nones K, Cowin P, Alsop K, Bailey PJ. 2015a. Whole–genome characterization of chemoresistant ovarian cancer. Nature 521:489–494.

Patch A-M, Christie EL, Etemadmoghadam D, Garsed DW, George J, Fereday S, Nones K, Cowin P, Alsop K, Bailey PJ et al. 2015b. Whole–genome characterization of chemoresistant ovarian cancer. Nature 521:489–494.

Rizzo S, Hersey JM, Mellor P, Dai W, Santos-Silva A, Liber Diea. 2011. Ovarian cancer stem cell-like side populations are enriched following chemotherapy and overexpress EZH2. Mol Cancer Ther 10:325–335.

Sellar GC, Watt KP, Rabiasz GJ, Stronach EA, Li L, Miller EP, Massie CE, Miller J, Contreras-Moreira B, Scott D. 2003. OPCML at 11q25 is epigenetically inactivated and has tumor-suppressor function in epithelial ovarian cancer. Nature genetics 34:337–343.

Vaughan S, Coward JI, Bast RC, Berchuck A, Berek JS, Brenton JD, Coukos G, Crum CC, Drapkin R, Etemadmoghadam D. 2011. Rethinking ovarian cancer: recommendations for improving outcomes. Nature Reviews Cancer 11:719–725.

Weber M, Hellmann I, Stadler MB, Ramos L, Pääbo S, Rebhan M, Schübeler D. 2007. Distribution, silencing potential and evolutionary impact of promoter DNA methylation in the human genome. Nature Genetics 39:457.

Yin Y, Morgunova E, Jolma A, Kaasinen E, Sahu B, Khund-Sayeed S, Das PK, Kivioja T, Dave K, Zhong F. 2017. Impact of cytosine methylation on DNA binding specificities of human transcription factors. Science 356:eaaj2239.

Zeller C, Dai W, Steele N, Siddiq A, Walley A, Wilhelm-Benartzi C, Rizzo S, van der Zee A, Plumb J, Brown R. 2012. Candidate DNA methylation drivers of acquired cisplatin resistance in ovarian cancer identified by methylome and expression profiling. Oncogene 31:4567–4576.

