## Supplementary Figures S1-S6, Supplementary Table S1 for "Amplification of genomic regions harbouring genes with dose-limiting effects provides selection pressure for acquiring aberrant epigenetic silencing in ovarian cancer"

**Supplementary Figure S1 – Promoter CpG methylation tends to be low for the analysed loci**

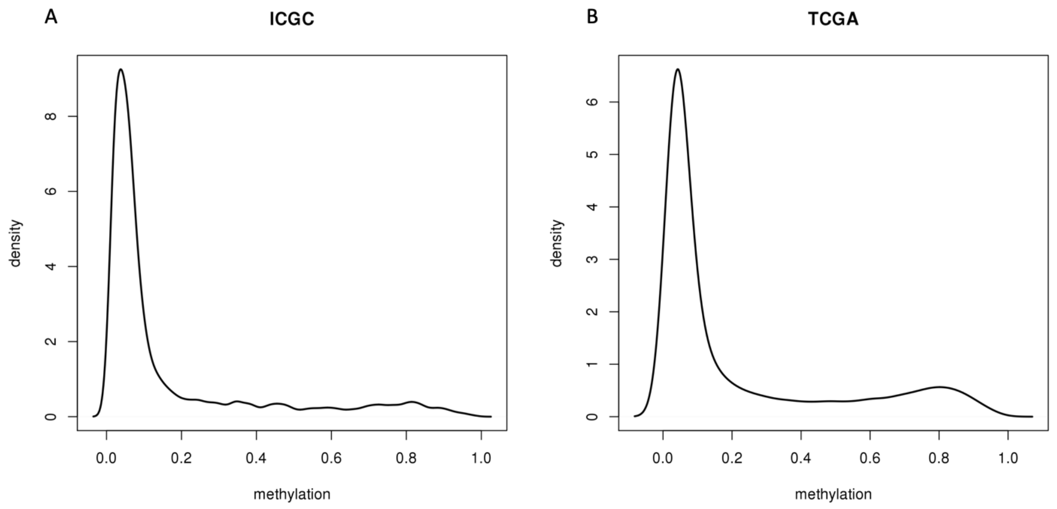


A) Kernel density estimation showing relative abundance of different levels of methylation (β-values) across 6680 CpG sites mapping to loci with at least 10% of samples in the ICGC cohort affected by copy-gain

B) As Fig.S1A but for 5084 CpG sites with at least 10% of samples in TCGA cohort affected by copy-gain

**Supplementary Figure S2 – Additional genome-wide analyses of correlation between copy-number and DNA methylation**

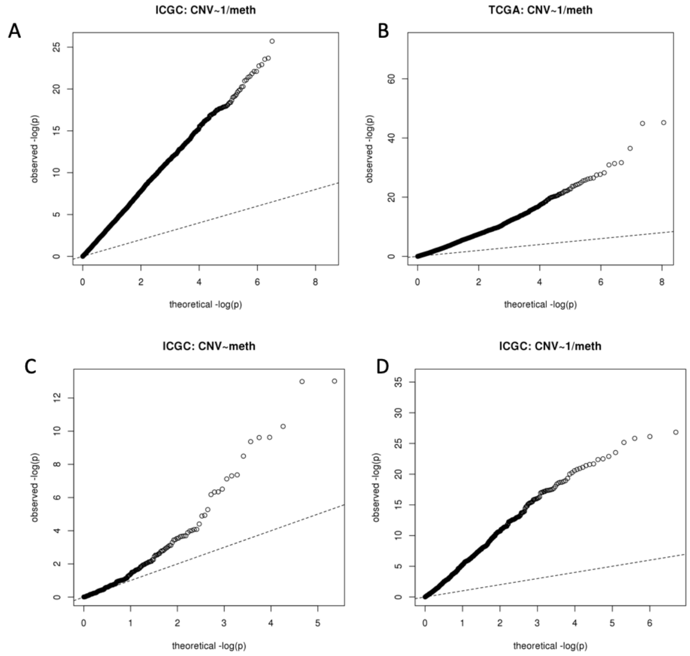


A) Quantile-quantile plot showing the distribution of significance estimates (-log p-values) of negative Spearman correlation coefficients between gCN and DNA methylation in the ICGC dataset, compared to the theoretical distribution under independence of gCN and DNA methylation for all loci (uniformly-distributed p-values, indicated by the dashed line).

B) As Fig.S2A but for TCGA cohort.

C) Quantile-quantile plot showing the distribution of significance estimates (-log p-values) of Spearman partial correlation coefficients between gCN and average DNA methylation across a CpG island in the ICGC dataset, controlling for cancer cell content of the corresponding tumour samples. Dashed line gives the theoretical distribution under independence of gCN and DNA methylation, when adjusting for differences in cancer cell content, for all loci (uniformly-distributed p-values).

D) As Fig.S2C but for TCGA cohort.

**Supplementary Figure S3 – Illustrations of correlation between copy-number, DNA methylation and tumour cellularity**

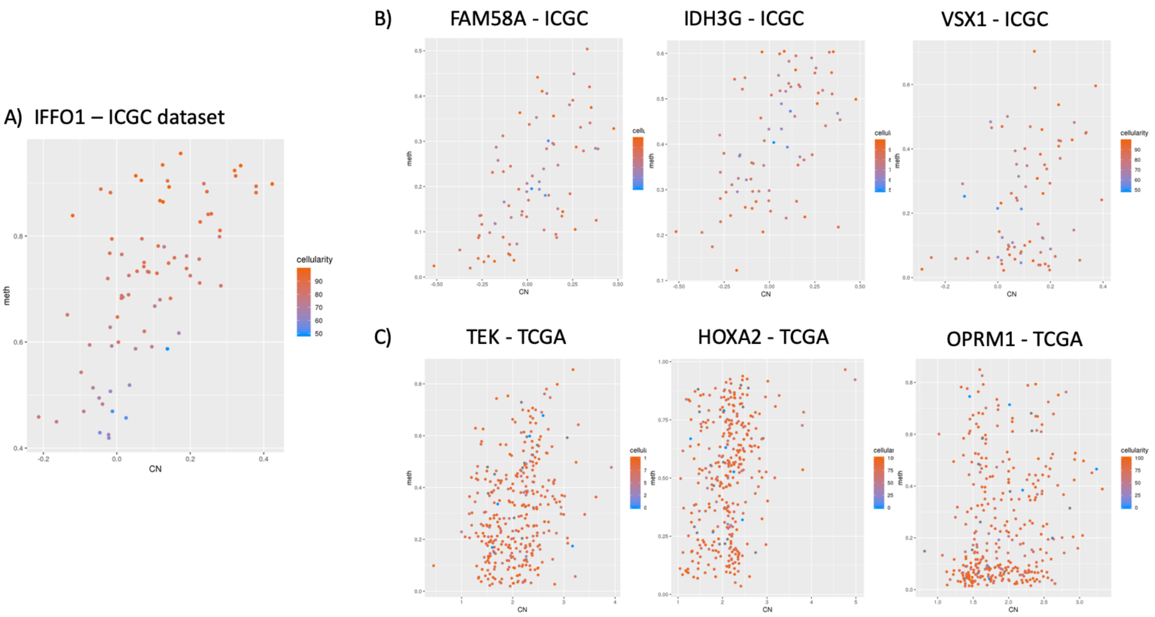


A) Plot of DNA methylation beta-value (y-axis) against copy-number (x-axis, as $log_2(CN/2)$) for 1 CpG locus (cg23737737) in the IFFO1 promoter, for tumour samples in the ICGC cohort. Points are coloured by tumour cellularity (percentage cancer cell content in tissue sample).
B) Plot of DNA methylation beta-value (y-axis) against copy-number (x-axis, as $log_2(CN/2)$) for 3 CpG loci (cg06511189, cg27271445, cg01715455) in the promoters of FAM58A, IDH3G and VSX1 (respectively), for tumour samples in the ICGC cohort. Points are coloured by tumour cellularity (percentage cancer cell content in tissue sample).
C) Plot of DNA methylation beta-value (y-axis) against copy-number (x-axis) for 3 CpG loci (cg09827833, cg26069745, cg14262937) in the promoters of TEK, HOXA2 and OPRM1 (respectively), for tumour samples in the ICGC cohort. Points are coloured by tumour cellularity (percentage cancer cell content in tissue sample).

### Supplementary Figure S4 - Compensatory DNA methylation in regions of genomic copy-gain or loss

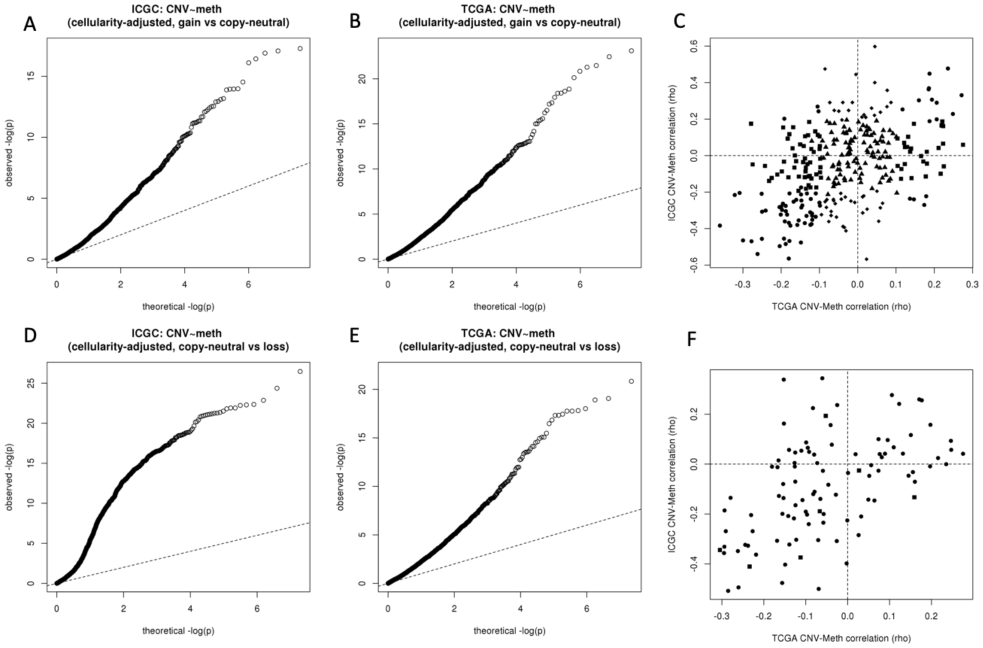


A) Quantile-quantile plot showing the distribution of significance estimates (-log p-values) of Spearman partial correlation coefficients between gCN and DNA methylation in the ICGC dataset, controlling for cancer cell content of the corresponding tumour samples and excluding samples affected by copy-loss at the corresponding locus. Dashed line gives the theoretical distribution under independence of gCN and DNA methylation, when adjusting for differences in cancer cell content, for all loci (uniformly-distributed p-values).
B) As Fig.1A but for TCGA cohort.
C) Plot illustrating conservation of locus-wise cellularity-adjusted gCN to DNA methylation partial correlations across the two HGSOC cohorts analysed, having excluded samples affected by copy-loss at the corresponding locus. X-axis shows partial Spearman correlation coefficient relating gCN to methylation (adjusted for cancer cell content) in TCGA cohort, y-axis shows partial Spearman correlation coefficient relating gCN to methylation (adjusted for cancer cell content) in ICGC cohort.
A) Quantile-quantile plot showing the distribution of significance estimates (-log p-values) of Spearman partial correlation coefficients between gCN and DNA methylation in the ICGC dataset, controlling for cancer cell content of the corresponding tumour samples and excluding samples affected by copy-gain at the corresponding locus. Dashed line gives the theoretical distribution under independence of gCN and DNA methylation, when adjusting for differences in cancer cell content, for all loci (uniformly-distributed p-values).
B) As Fig.1A but for TCGA cohort.
C) Plot illustrating conservation of locus-wise cellularity-adjusted gCN to DNA methylation partial correlations across the two HGSOC cohorts analysed, having excluded samples affected by copy-gain at the corresponding locus. X-axis shows partial Spearman correlation coefficient relating gCN to methylation (adjusted for cancer cell content) in TCGA cohort, y-axis shows partial Spearman correlation coefficient relating gCN to methylation (adjusted for cancer cell content) in ICGC cohort.

### Supplementary Figure S5 – Survival associations of DNA methylation at DCPM loci

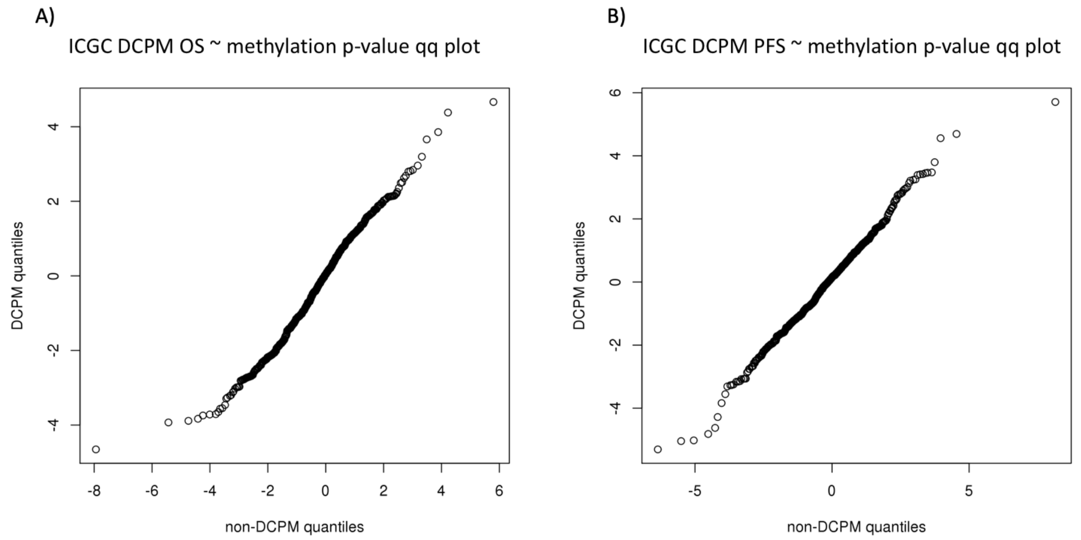


A) Quantile-quantile plot showing distribution of statistical significance (as -log(p-value)) of associations between DNA methylation β-values and patient overall survival in the ICGC cohort. Statistical significance was estimated through fitting Cox proportional hazards regression model.

B) Quantile-quantile plot showing distribution of statistical significance (as -log(p-value)) of associations between DNA methylation β-values and patient progression-free survival in the ICGC cohort. Statistical significance was estimated through fitting Cox proportional hazards regression model.

**Supplementary Figure S6 – Correlation of DNA methylation of individual CpG sites within a CpG island**

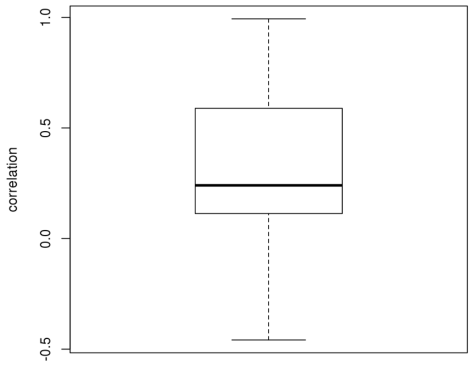


Box plot showing distribution of Pearson correlation coefficients computed between the methylation β-values for each pair of CpG sites sharing a mapping to the same CpG island, in the ICGC cohort.

**Supplementary Table S1 – Annotation of CpG sites affected by DCPM**

**Supplementary Table S2 – Complete data for RT-PCR assays**

**Supplementary Table S2 – Complete data for pyrosequencing assays**

**Supplementary Table S4 – Primers for RT-PCR and pyrosequencing**

#### Primers for RT-PCR

Primers designed, verified with *in silico PCR* (*UCSC Genome Browser*) and obtained from Sigma-Aldrich to verify gene expression using RT-PCR (diluted to 5μM). Optimised through a primer test – standard curve.

| Gene | *Forward primer* | *Reverse primer* |
|---|---|---|
| ZMAT3 | GCTCTGTGATGCCTCCTTCAGT | TTGACCCAGCTCTGAGGATTCC |
| ECT2 | GCAGTCAGCAAGGTGGCAAGTT | CTCTGGTGCAAGGATAGGTCCA |
| PIK3CA | GAAGCACCTGAATAGGCAAGTCG | GAGCATCCATGAAATCTGGTCGC |
| GPR171 | AGCCAAAGAGGCTACACTGCTC | CCTTGGTCTCTTTAGGTGAGGC |
| SIAH2 | GCATCAGGAACCTGGCTATGGA | GCAGGAGTAGGGACGGTATTCA |
| RAP2B | CGCAAGGAGATTGAGGTGGACT | TTGACGAGGCTGTAGACCAGGA |

#### Primers for pyrosequencing

| Gene - locus | *Forward primer* | *Reverse primer* |
|---|---|---|
| ZMAT3 - cg07266910 | TTGAGGAGAAGGAGGAAATTTT | AACCTCTTAACATAATTCTTCCCTTAATA |
| GPR171 - cg11630392 | GTTAGTAGTTTGTAATGGGGTTGT | CCTAAAACTATCTTCTATAAACCTCAAC |
